# Relative humidity differentially modulates decay of structurally distinct respiratory viruses in deposited human saliva particles

**DOI:** 10.64898/2026.09.23.753914

**Authors:** Margot Olive, Zachariah Broemmel, Sterling Knight, Nicole C. Rockey

## Abstract

Respiratory viruses pose a major threat to human health. Environmental virus persistence in respiratory secretions is essential for successful spread, yet studies have focused largely on a select few viruses in non-physiological matrices. Here, we compared the stability of structurally distinct human respiratory viruses, adenovirus, influenza virus, and rhinovirus, in deposited particles of pooled human saliva under indoor-relevant relative humidity (RH) conditions. Adenovirus exhibited <2.2-log_10_ decay over 6 h regardless of RH, while rhinovirus exhibited significantly greater degradation (>3.8-log_10_, at 50% and 80% RH). Influenza virus showed minimal decay at 20% and 50% RH but degraded significantly more (3-log_10_) at 80% RH. To distinguish aspects of human saliva that are key determinants in driving virus decay, we recapitulated saliva composition using the inorganic solute and protein concentrations determined via ion-coupled plasma mass spectrometry and proteomics, respectively. Rhinovirus was significantly more stable in simulated saliva than in human saliva at 50% RH, while influenza virus and adenovirus behaved similarly in both matrices. These findings emphasize virus-specific environmental persistence and highlight limitations of relying on a single virus and particle matrix for defining public health strategies. Ultimately, these insights will contribute to approaches for mitigating environmental transmission of distinct respiratory pathogens.

## Introduction

Human respiratory viruses spread through the environment following emission of virus-laden particles, either deposited or airborne, from an infected individual.^1,2^ Thus, successful transmission is predicated on a virus’ ability to remain infectious outside the host. Understanding environmental persistence of the human respiratory viruses that cause disease year over year is therefore essential for accurately defining transmission risks and designing effective mitigation strategies to combat spread.

Virus susceptibility outside the host is governed by numerous factors, including virus structure and type, virus microenvironment, temperature, and relative humidity (RH).^3^ Viruses exhibit broad diversity in their size, morphology, presence or absence of a lipid membrane, protein makeup, and nucleic acid type, among other differences.^4^ Given their disparate structural attributes, it is not surprising that viruses can be differentially susceptible to environmental exposures outside the host. Indeed, enveloped viruses, like influenza virus, respiratory syncytial virus (RSV), and severe acute respiratory syndrome coronavirus 2 (SARS-CoV-2), have generally been shown to exhibit enhanced stability at low RH, with elevated decay at midrange RH,^5–9^ while trends for nonenveloped viruses vary depending on the virus and the deposited or airborne nature of the virus suspension.^10–12^ Aerosol studies of nonenveloped picornaviruses including poliovirus, for example, commonly observed rapid decay at low RH, with improved stability at elevated RH,^11,13–20^ while inconsistent effects of degradation on surfaces with variable RH have been observed.^10,12,21,22^

While the persistence of a select few respiratory viruses has been well-studied in environmentally relevant transmission scenarios,^3,5–7,7–9,11,14,23–30^ work to date has largely overlooked a portion of respiratory viruses that spread in human populations, particularly those with viral structures distinct from the enveloped RNA viruses influenza virus and coronavirus. This is despite the fact that nonenveloped respiratory viruses, like adenovirus and rhinovirus, contribute considerably to global disease burden and circulate regularly.^31–33^ Rhinoviruses, for example, are the most common cause of the cold worldwide, accounting for ∼13–59% of detections globally and up to ∼60% of acute respiratory infections in young children.^34^

A limited number of studies have explored the environmental susceptibility of adenovirus in aerosols, demonstrating rapid decay at low and midrange RH and minimal decay at high RH over the same timeframe.^20^ Buckland and Tyrell observed a similar trend on surfaces, with 1 to 2-log_10_ decay of three adenovirus strains at low RH after 2.5 hrs of exposure, and only 0 to 1-log_10_ decay at high RH.^10^ In contrast, RH had no impact on the degradation of an enteric adenovirus on surfaces,^21^ and an ophthalmology study detected infectious adenovirus for up to 11 days following inoculation on medical tool surfaces at room temperature, though RH was not reported.^35^ Research with human rhinovirus also shows mixed findings, with rhinovirus exhibiting enhanced decay at low RH but retaining infectivity at elevated RH,^36^ while Niazi et al. observed a V-shaped relationship in airborne HRV16 persistence with RH.^37^ Low levels of rhinovirus degradation were observed on surfaces at both low and high RH.^10^ The small number of nonenveloped respiratory virus studies conducted under a narrow set of conditions leaves open questions about how these viruses persist in environmental settings relevant for virus transmission. A better understanding of the role virus structural attributes play in driving stability outside the host is critical to improving the effectiveness of interventions targeted at reducing virus spread.

The virus microenvironment, including the constituents present within the emitted matrix encasing the virion, also influences virus susceptibility.^13,29,38,39^ Emitted respiratory matrices typically include nasal mucus, respiratory mucus, or saliva, and these physiological solutions predominantly consist of water (>95%), although a complex range of macromolecules, including proteins, lipids, and salts in the ranges of ∼100 to 1000 mg/dL, ∼1 to 1000 mg/dL, and ∼0.5 to 10,000 mg/dL, respectively, is typical.^40,41^ The presence of concentrated salts in airborne or deposited particles rapidly inactivates virions,^13,42^ while concentrated proteins in these particles can stabilize viruses and enhance persistence.^5,23,43^ Proteins such as fetal bovine serum, bovine serum albumin, and porcine mucin have been shown to stabilize viruses in airborne and deposited particles at variable RH.^5,23,26,43^ Bulk protein concentration alone does not drive virus decay or persistence, indicating the importance of protein type in establishing stability.^6,23^ Particle crystallization and colocalization of viruses with proteins or crystals in dried particles have been proposed as potential mechanisms for virus decay or protection.^6,26,44,45^ Interestingly, work by Schaub et al. indicates that salt molality, rather than salt crystallization and particle drying morphology itself, is a driver of degradation.^42^ While these findings provide much-needed mechanistic insights into virus decay, they primarily rely on the use of laboratory-derived matrices comprised of physiologically imprecise constituents and concentrations that limit their applicability to real-world virus persistence and subsequent exposure risks. Improved understanding of virus decay in physiologically accurate solutions is needed. In addition, research to determine if simplistic solutions can correctly recapitulate virus decay in real respiratory matrices is critical to advancing the standardization and pertinence of environmental virus persistence work.

In this study, we quantified the persistence of three structurally distinct respiratory viruses, namely human adenovirus type 4 (HAdV4), human rhinovirus A16 (HRV16), and influenza A virus A/CA/07/2009 (H1N1pdm09) (Supporting Information (SI) Table S1), in deposited particles comprised of pooled human saliva under controlled environmental conditions (i.e., ambient temperature and 20%, 50%, and 80% RH) representative of indoor spaces, where respiratory virus transmission risks are elevated.^46^ Further, we characterized the pooled human saliva and used this information to generate a simulated saliva that we could use to evaluate if our simplified saliva solution recapitulated the observed virus decay trends in human saliva. By directly comparing three distinct human viruses in physiologically relevant fluids, this study provides critical data on virus-specific persistence, informing risk assessments and guiding strategies to reduce respiratory virus transmission in indoor environments.

## Materials and Methods

### Cell maintenance

All cell lines were maintained in monolayer cultures at 37°C and 5% CO_2_. MDCK cells were maintained in Eagles’ Minimum Essential Medium (MEM; Millipore Sigma, Cat. No. M2279) supplemented with 10% Serum Plus II (Millipore Sigma, Cat. No. 14009C), 1% penicillin-streptomycin (Thermo Fisher Scientific, Cat. No. 15140122), and 1% L-glutamine (Thermo Fisher Scientific, Cat. No. 25030081). A549 and H1HeLa cells were maintained in Dulbecco’s Modified Eagle Medium (DMEM; Thermo Fisher Scientific, Cat. No. 11965092) containing high glucose and supplemented with 10% Serum Plus II, 1% penicillin-streptomycin, and 1% L-glutamine.

### Virus propagation and stock preparation

Three human respiratory viruses were used in this study. Specifically, HAdV4 (American Type Culture Collection (ATCC), Cat. No. VR-1572), HRV16, kindly provided by Dr. James Gern, and H1N1pdm09, kindly provided by Dr. Seema Lakdawala, were used. The selected viruses differ in virion structure, including dissimilarities in size, genome type and length, and the presence or absence of a lipid envelope (Table S1). Details of virus propagation and stock preparation are provided in the SI, including details of the pooled human saliva (Table S2) and simulated saliva matrices used to resuspend virus stocks. Infectious virus concentrations in pooled human saliva and simulated saliva stocks are available in Table S3.

### Bulk human saliva and virus stock characterization

To control for key microenvironment factors that could influence virus persistence, we measured bulk protein concentrations, sucrose concentrations, elemental concentrations, conductivity, and pH in virus-free pooled human saliva, virus-free protein-salt matrix and in pooled human saliva virus stocks. Bulk protein concentrations were determined using the Pierce bicinchoninic acid (BCA) assay (Thermo Fisher Scientific, Cat No. 23227) with bovine serum albumin (BSA) as the standard. Sucrose was measured using a modified sucrose assay (Sigma Aldrich, Cat No. SCA20-1KT) following manufacturer’s protocol - except that microplates were used instead of cuvettes - and the absorbance was measured at 340 nm with a spectrophotometer (Molecular devices, SpectraMax iD3 microplate reader). At least two independent replicates were conducted for sucrose and BCA assays.

Elemental concentrations in pooled human saliva were quantified using inductively coupled-plasma mass spectrometry (ICP-MS) in two independent replicates as described previously.^47^ Briefly, 200 µL samples were digested with 500 µL nitric acid and heated at 95°C for 30 minutes. Samples were then cooled to room temperature, followed by addition of 500 µL hydrogen peroxide and heating at 95°C for 30 min. To compensate for evaporation during heating, 300 µL milliQ was added to each sample following heating. 100 µL sample digestate was subsequently analyzed by ICP-MS for Na, K, Ca, Mg, and Al. Raw data were corrected to account for sample volume dilution and determine elemental concentrations in samples. Conductivity and pH were quantified using an InLab 731 conductivity probe (Mettler Toledo, Cat. No. MT30014092) and InLab Expert Pro-ISM electrode (Mettler Toledo, Cat. No. MT30014096), respectively, connected to a Multiparameter SevenExcellence Meter (Mettler Toledo).

### Human saliva protein profile

We characterized the proteins present in our purchased pooled human saliva via liquid chromatograph coupled to tandem mass spectrometry (LC-MS/MS) at the Duke Proteomics and Metabolomics Core Facility. Saliva proteins were denatured, reduced, alkylated, and digested with trypsin, and the resulting peptides were analyzed using an Evosep One LC interfaced to a ThermoFisher Orbitrap Astral. Details of proteomics sample preparation, data acquisition, and downstream analysis are provided in the SI.

### Deposited particle generation, exposure, and collection

10 x 1 μL deposited particles of each virus stock in pooled human saliva and simulated saliva suspensions were deposited on sterile, non-treated, polystyrene 6-well plates (Costar, Cat. No. 3736) and exposed to typical indoor temperature (22.5°C) and variable RH (i.e., 20%, 50% or 80%) for up to 6 h in a temperature and humidity controlled environmental chamber (Plas Labs, Model 810-SG10). A Hobo Temperature and Humidity Data logger (Onset, Cat. No. UX100-011A) was used to monitor temperature and RH during experiments (Figures S1 and S2). For virus-laden human saliva suspensions, deposited particles were collected following 0 min, 15 min, 30 min, 60 min, 120 min, 240 min, or 360 min of environmental exposure (and up to 24 hours for HAdV4). For virus-laden simulated saliva suspensions, deposited particles were exposed to 50% RH and collected following 0 min, 60 min, 120 min, or 240 min of environmental exposure. After exposure, deposited particles containing H1N1pdm09 were resuspended in 500 μL of MEM supplemented with 1% L-glutamine, and deposited particles containing HAdV4 and HRV16 were resuspended in 500 µL of DMEM containing L-glutamine.

Following collection, samples were immediately stored at −80°C until downstream analyses. At least three independent replicates were performed for each RH condition and deposited particle composition, with technical duplicates included for every independent replicate. Bulk controls of the deposited particle solutions were collected at 0 min and post-experimentation by diluting 10 µL of the bulk solution into 500 µL of the corresponding medium to quantify any virus decay attributable to bulk-phase effects.

### Nucleic acid recovery after deposited particle drying

Virus recovery was quantified by measuring nucleic acid levels in virus-laden human saliva particle samples for a subset of 50% RH exposures (i.e., 0 min, 30 min, 120 min, and 360 min) to ensure collection processes did not contribute to reduced infectious virus concentrations following exposure experiments. Two independent replicates of each deposited particle virus decay experiment were assessed for virus recovery. Additional assay details, including extraction and (RT)-qPCR procedures, primer and probe sequences, cycling conditions, amplicon targets, and standard generation, are included in the SI text and Table S4.

### Particle drying

Virus-laden deposited particle drying times were determined by recording particle drying during environmental exposures. The same deposited particle generation process was used as described above. Videos were acquired until drying of all deposited particles at 20%, 50%, and 80% RH for all three viruses in the pooled human saliva solution, while videos were conducted for all three viruses in the simulated saliva solution at 50% RH. Drying time was determined by recording the times associated with the first and last deposited particle drying in a set of 10 x 1 µL deposited particles. Reported drying times are the average of two independent replicates. At the end of each drying experiment, the morphology of at least three deposited particles was captured using a bright field microscope (Olympus, Cat. No. CKX53SF) equipped with an Olympus EP50 camera. All images were acquired at 4x magnification using EpiView software (version 1.4).

### Virus decay kinetics

Given that a range of models has been used to describe the kinetics of virus decay in deposited particles, ^3,6,27,48^ we fit our data to two candidate models to determine which best represented our experimental data. These models included a single-phase first order decay model and a two-phase first order decay model. The single-phase first order decay model was defined as:

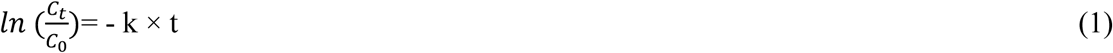

Where C and C_t_ represent the infectious virus concentrations in deposited particles at time 0 and time t, in minutes, respectively, and k represents the first-order decay rate constant in min^-1^. The two-phase decay model consisted of two first-order decay phases separated by a breakpoint t = t_dry_ corresponding to the time at which the deposited particles fully dried based on video analysis, described as follows.

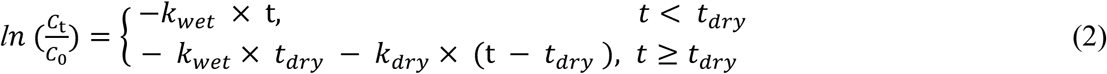

Where C and C_t_ represent the infectious virus concentrations in deposited particles at time 0 and time t, in minutes, respectively, and *k_wet_* and *k_dry_* represent the first-order decay rate constant in min^-1^ during the wet and dry phases, respectively.

### Statistical analysis

All statistical analyses were performed in GraphPad Prism 9 (version 9.5.1).^49^ The two-phase decay model described in equation 2 corresponds to the "segmental linear regression" analysis on Prism. Statistical significance was defined at an α value of 0.05.

## Results

### HRV16 decays rapidly at elevated RH in deposited human saliva particles, while HAdV4 persists similarly regardless of RH

We used a tightly controlled temperature and humidity deposited particle system to investigate the persistence of structurally distinct respiratory viruses in human saliva following exposure to variable RH (i.e., 20%, 50%, and 80% RH) for up to six hours. All three viruses were most stable at 20% RH, with 0.7-log_10_ decay, 0.5-log_10_ decay, and 1.8-log_10_ decay, on average, for HAdV4, H1N1pdm09, and HRV16, respectively, at 6 h post-exposure (Figure 1). HAdV4 maintained infectivity across all RHs for up to six hours of exposure, with less than 2.2-log_10_ decay at each RH condition, while H1N1pdm09 and HRV16 exhibited intermediate decay at 50% RH and the greatest decay at 80% RH (Figure 1). No statistically significant differences in infectivity loss across RH conditions were observed for HAdV4 after four hours of exposure (all p > 0.05, ANOVA; Table S5). In contrast, significant differences in H1N1pdm09 infectivity loss between 20% and 80% RH were evident beyond 2 h of exposure (all p < 0.05, ANOVA; Table S5), and HRV16 showed significantly more decay at 80% compared to at 20% RH at 1h, 4h and 6 h post-exposure (all p < 0.05, ANOVA; Table S5). Due to its elevated persistence, we also assessed HAdV4 stability at 24 hours of exposure to 50% RH, and HAdV4 exhibited insignificant levels of additional decay over this period, with < 3-log_10_ total decay (p > 0.05, unpaired two-tailed t-test; Figure S3).

**Figure 1.**
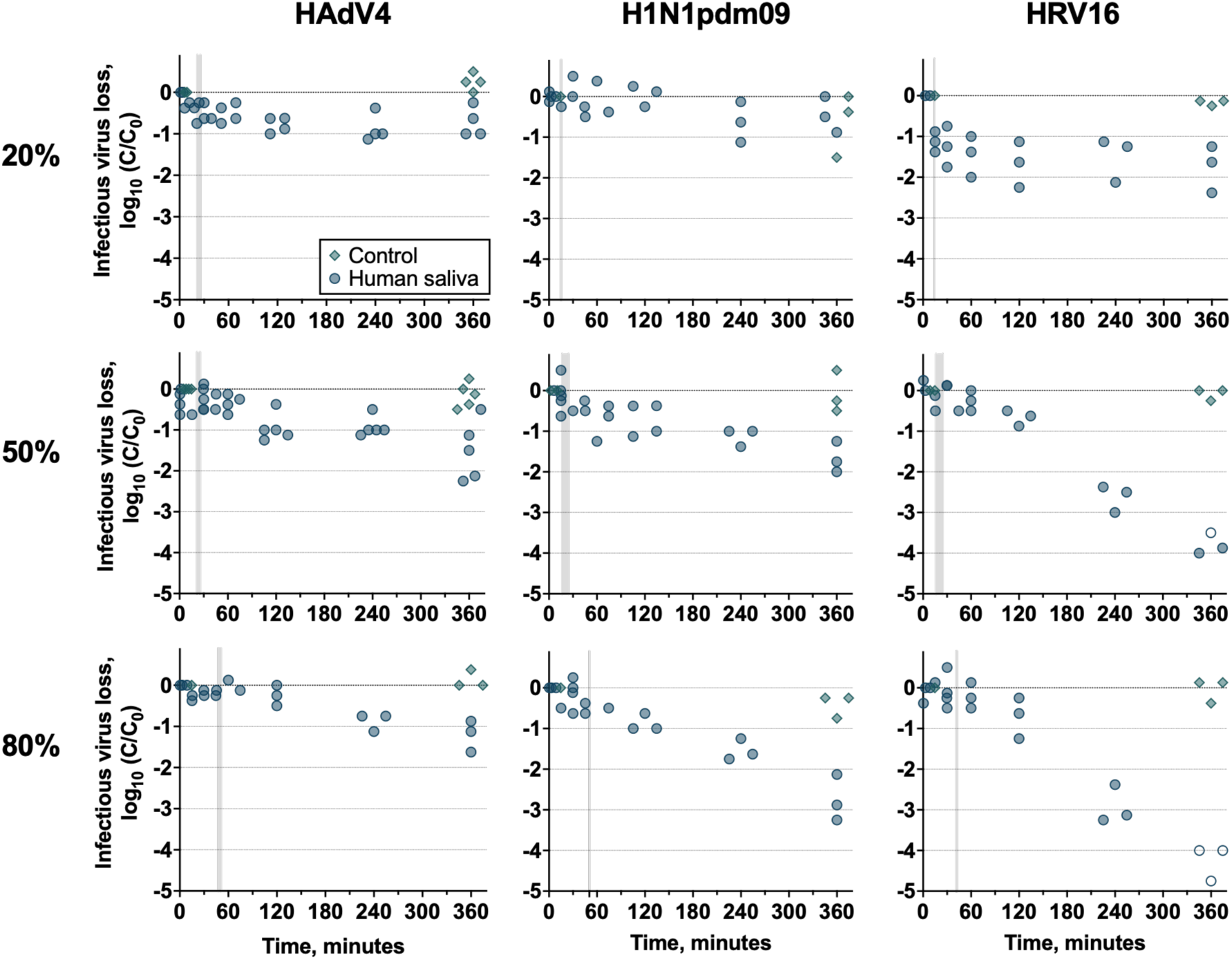
Decay of infectious HAdV4, H1N1pdm09, and HRV16 in deposited human saliva particles exposed to 20%, 50%, or 80% RH and 22.5°C. Two technical replicates were conducted for each independent replicate, and the mean decay of two technical replicates is shown for each independent replicate (n ≥ 3). Empty symbols represent data below detection. Gray shading represents deposited particle drying time (Table S7). Controls are bulk solution samples taken at the beginning and end of each experiment.

While HRV16 was more persistent at 20% RH than at higher RHs, this virus exhibited significantly more decay than H1N1pdm09 beyond 1 h of exposure (all p < 0.05, ANOVA; Table S6). Following 6 h at 50% RH, HRV16 infectivity in deposited saliva particles decayed significantly more than HAdV4 and H1N1pdm09 under the same conditions, with > 4.0-log_10_ decay compared to 2.3-log_10_ and 2.0-log_10_ decay for HAdV4 and H1N1pdm09, respectively (all p < 0.05; ANOVA; Table S6). H1N1pdm09 and HAdV4 infectivity losses were not significantly different at 50% RH (all p > 0.05, ANOVA; Table S6). Trends were similar at 80% RH, with HAdV4 and H1N1pdm09 exhibiting significantly more stability than HRV16 in deposited saliva particles at 4 h of exposure and beyond (all p < 0.05 for HAdV4 vs HRV16; all p < 0.05 for H1N1pdm09 vs HRV16, ANOVA; Table S6), and no significant differences in HAdV4 and H1N1pdm09 decay at 4 h (p > 0.05, ANOVA; Table S6).

Nucleic acid recovery was conducted for a subset of experiments at 50% RH for all three viruses to ensure that any decay we observed in infectious virus was not due to poor recovery of viruses from deposited particles. Recoveries at 0.5 h, 2 h, and 6 h were not significantly different from recovery at 0 h for any of the viruses (all p > 0.05, ANOVA; Figure S4). Together, these results confirm that the losses of infectivity measured over the course of our experiments were not due to virus losses from deposited particle recovery.

### Rapid virus decay was best described using a single-phase linear model, while a two-phase model was more suitable for minimal virus decay

A two-phase decay curve has been used to accurately describe the kinetics of infectious virus degradation in deposited particles during environmental exposure.^3,6,48^ In this model, the initial curve is associated with the drying particle phase, and the subsequent curve is associated with the dried or equilibrium particle phase.^6,48^ We determined particle drying times following deposition to identify the time point that delineated these two phases of decay. As expected, particles took more time to dry as RH increased, regardless of the virus (Table S7). We evaluated all decay curves using single phase and two-phase linear models and assessed goodness of fit to determine which model best described decay kinetics (Figure S5, Table S8).

Interestingly, for virus decay curves showing pronounced decay, specifically that of HRV16 at 50% and 80% RH and H1N1pdm09 at 80% RH, decay kinetics were described similarly by a single linear model and a two-phase model, with goodness-of-fit values exceeding 0.91 regardless of model type. On the contrary, decay curves showing a drop in infectivity followed by a plateau, namely the curves for HAdV4 at 20% RH and more notably HRV16 at 20% RH, were not well described by a single-phase linear model. By applying a two-phase decay curve, the trends could be better explained and resulted in a moderate fit with an average goodness of fit of 0.6.

For decay curves in which the two model types performed significantly differently, we selected the model type that performed best, while for decay curves that did not result in significant differences across the two model types, we used a single-phase linear model for simplicity. From these models, we determined the rate constants for each virus at 20%, 50%, and 80% RH (Table S8). Across the RH conditions tested, the decay rate constants for HAdV4 were low. At 20% RH, the wet-phase rate constant was 0.032 ± 0.018 min^-1^ (slope ± 95% CI), while the dry-phase rate constant was minimal (0.001 ± 0.002 min^-1^) but remained significantly different from zero (p *<* 0.05, F-test; Table S9). At 50% and 80% RH, HAdV4 decay rates were low, with rate constants of 0004 ± 0.001 min^-1^ and 0.003 ± 0.001 min^-1^, respectively, and were not significantly different from one another (p > 0.05, F-test; Table S9).

In contrast, the decay rate constants for H1N1pdm09 and HRV16 varied significantly with RH. H1N1pdm09 showed minimal decay kinetics at 20% RH (k = 0.002 ± 0.001 min^-1^) but increased significantly at 50% RH (k = 0.004 ± 0.001 min^-1^; p < 0.05, F-test). Relative to 50% RH, the rate constant for H1N1pdm09 decay at 80% RH was even higher (k = 0.007 ± 0.001 min^-1^; p < 0.05, F-test). HRV16 decayed comparably at 50% and 80% RH (k = 0.011 ± 0.001 min^-1^ and k = 0.012 ± 0.002 min^-1^, respectively; p > 0.05, F-test), however at 20% RH, kinetics exhibited a two-phase pattern, with rapid decay during the wet phase (k_wet_ = 0.092 ± 0.043 min^-1^) followed by negligible decay during the dry phase (k_dry_ = 0.001 ± 0.001 min^-1^; p > 0.05, F-test, Table S9).

When comparing viruses within each RH condition, all viruses showed slow post-drying decay at 20% RH with rate constants not exceeding 0.002 ± 0.001 min^-1^. At 50% RH, HRV16 decayed significantly faster (k = 0.011 h^-1^ ± 0.001 min^-1^) than either HAdV4 or H1N1pdm09 (both 0.004 ± 0.001 min^-1^; all p < 0.05, F-test). At 80% RH, both HRV16 (k = 0.012 ± 0.002 min^-1^) and H1N1pdm09 (k = 0.007 ± 0.001 min^-1^) decayed significantly faster than HAdV4 (k = 0.003 ± 0.001 min^-1^, p < 0.05, F-test).

### Pooled human saliva contains a diverse proteome with abundant alpha amylase, albumin, immunoglobulin, and established antiviral compounds

The stability of infectious viruses in deposited particles is impacted by several microenvironmental parameters, with particle composition playing an important role. To identify the constituents of our virus-laden particles that could contribute to the observed trends in virus susceptibility, we characterized the inorganics and proteins present in our pooled human saliva. The total protein concentration of the saliva, as measured by BCA, was 730.04 ± 80.30 mg/L (Figure 2), similar to reported levels for saliva.^40^ Proteomics of our pooled human saliva revealed the abundance of nearly 2,000 distinct protein groups (Figure 2F, Table S10). The alpha amylase 1A, 1B, and 1C protein group, which contains amylases that catalyze the hydrolysis of starches into simple sugars,^50^ was most abundant, at 7.29-log_10_ abundance (Figure 2D), followed by albumin (6.91-log_10_ abundance). Among the top 20 proteins detected, several known proteins with antiviral properties were found, including mucin 5B and deleted in malignant brain tumors 1 protein (DMBT1).^6,51,52^ Immunoglobulin (Ig) protein groups were also frequently detected, with those belonging to the IgA2 and IgA1 classes being the third and fourth most abundant protein groups identified, respectively; this is not surprising, given that IgA is a mucosal Ig.^53^ Our saliva was also highly abundant in cystatin S and cystatin SN, a class of proteins known to inhibit viral proteases, such as cysteine protease.^54,55^

**Figure 2.**
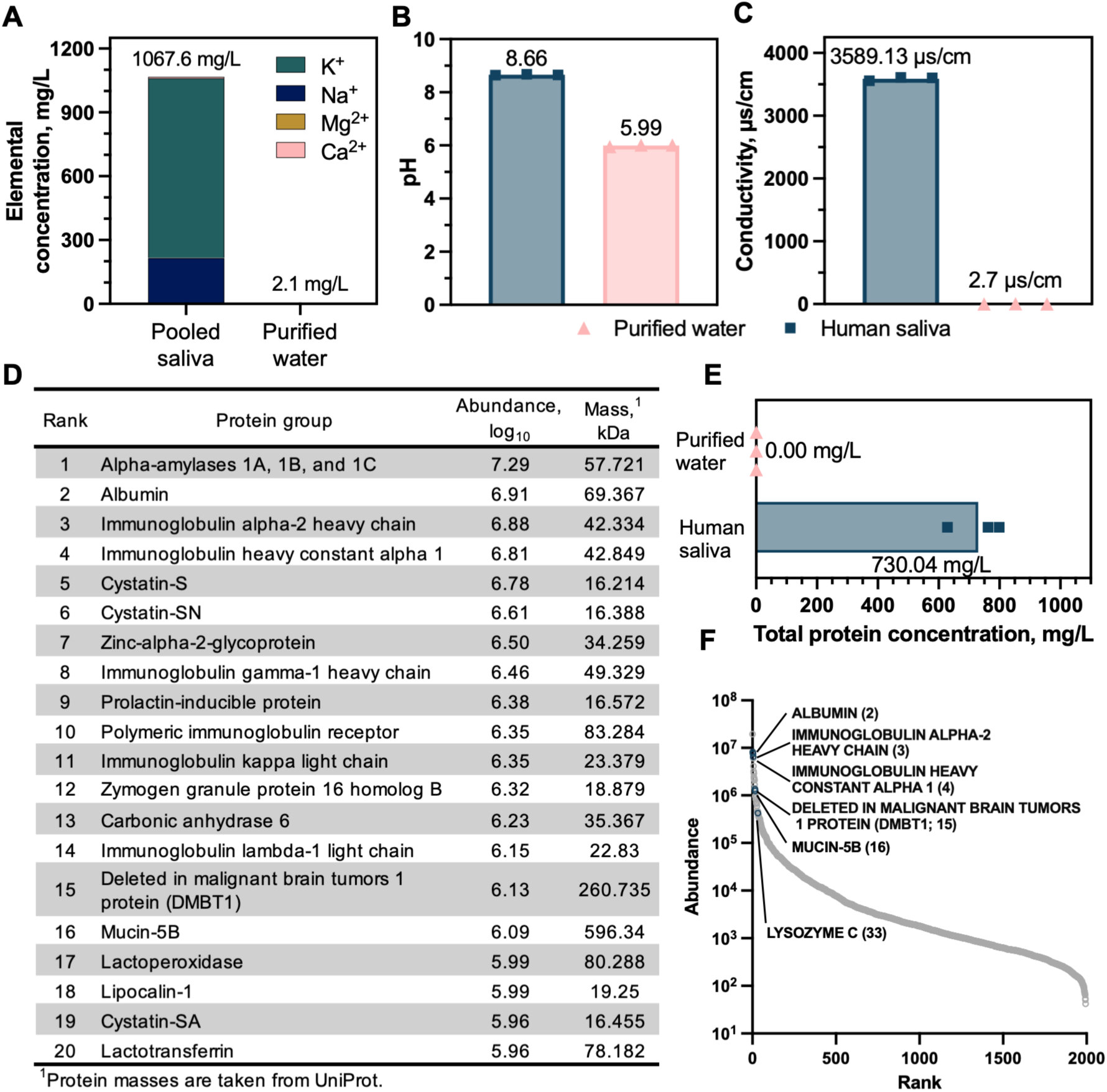
Characterization of pooled human saliva. (A) Elemental concentrations, (B) pH, and (C) conductivity of pooled human saliva in comparison to purified water. (D) 20 most abundant protein groups present in pooled human saliva and the associated abundance (log_10_) and mass of each group, as included in UniProt.^57^ (E) Total protein concentration in pooled human saliva as compared to purified water, and (F) rank abundance profile of protein groups identified via proteomics analysis. Highlighted protein groups in (F) are typical of human saliva or have been shown to play a role in virus inactivation. Rank abundance of the protein group is shown in parentheses. A complete table of protein group abundance is in Table S10.

To assess the inorganics in our saliva matrix, we measured elemental concentrations of K, Na, Mg, Ca, and Al via ICP-MS. K^+^ and Na^+^ were most abundant, with concentrations of 0.84 and 0.22 g/L, respectively (Figure 2A). Less concentrated cations included Ca^2+^ and Mg^2+^, with levels of 0.008 and 0.002 g/L, respectively, while Al was not detected. The total content of these elements was 1.067 g/L. Conductivity of the saliva was 3,589 µS/cm (Figure 2C) and pH was slightly basic, at 8.6 (Figure 2B). Since we resuspended our concentrated virus stocks in the pooled human saliva, and various aspects of virus propagation, purification, and resuspension can result in changes to virus stocks that may impact stability,^56^ we measured bulk protein, sucrose, and elemental concentrations of our virus stocks and confirmed that these constituents were not markedly different in our virus stocks compared to our pooled saliva itself (Table S11 and Figure 2).

### Decay of HAdV4 and H1N1pdm09 in simulated saliva particles recapitulates phenotypes observed in human saliva, but HRV16 degradation in simulated saliva does not

Ultimately, we sought to generate a simulated saliva matrix that could be used to recapitulate the same virus susceptibility kinetics we observed in human saliva. To achieve this, we added KCl, NaCl, CaCl_2_ and MgCl_2_ to sterile purified water at the same elemental concentrations detected via ICP-MS in our pooled human saliva specimen. With ∼ 2,000 different protein types in our human saliva, it was not feasible to completely recapitulate protein quantity and quality. We therefore simplified the simulated saliva protein composition by adding a single protein, human serum albumin, that was one of the most abundant proteins we identified via proteomics. Human serum albumin was added to the simulated saliva solution at the same total protein concentration, 730.04 ± 80.30 µg/mL, as was measured in human saliva. pH of the resulting simulated saliva was slightly acidic, 6.2 ± 0.08, as the purified water used had pH 5.99 ± 0.03 (Figure 2B). Virus stocks were resuspended in this simulated saliva and used to generate deposited particles exposed to 50% RH. HAdV4 and H1N1pdm09 persistence in our deposited simulated saliva particles was similar to stability in deposited human saliva particles (Figure 3, Table S12). HRV16, however, exhibited significantly less decay, 2.21-log_10_ less on average, in simulated saliva particles compared to in human saliva particles at 4 h of 50% RH exposure (p < 0.05, unpaired two-tailed t-test, Table S12). Bulk solution controls showed no significant decay (Figure S6).

**Figure 3.**
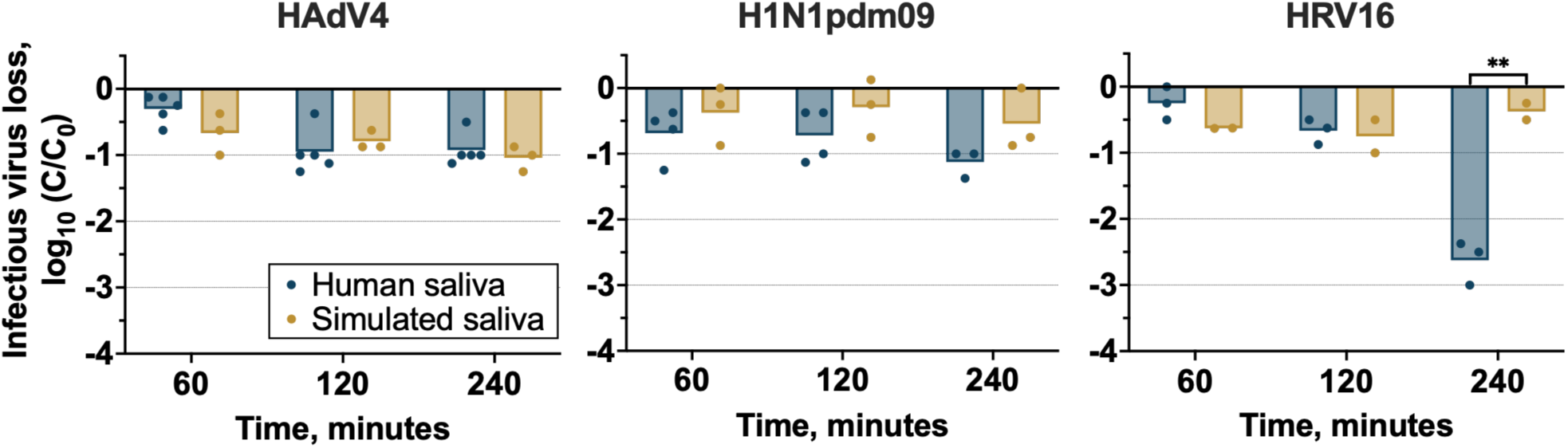
Decay of infectious HAdV4, H1N1pdm09, and HRV16 in pooled and simulated saliva matrices following exposure to 50% RH and ambient temperature for 60 min, 120 min or 240 min. Two technical replicates were conducted for each independent replicate, and the mean decay of two technical replicates is shown for each independent replicate (n ≥ 3). Vertical bars are the mean value of independent replicates. ** indicate a p-value < 0.005, unpaired t-test.

### Dried deposited particle morphology varies for H1N1pdm09 and HRV16 in saliva but not for HAdV4

Particle drying and the resulting interactions of virions with different particle constituents, like mucins or salts, have been hypothesized to explain environmental virus degradation in particles.^6,26,45,58,59^ We therefore investigated deposited virus-laden particles following drying under variable RH and particle composition to assess how deposited particle morphology correlated with our observed trends in virus decay. When evaluating morphology across RHs in dried human saliva particles, we observed elongated, extensive, thin crystalline structures at 20% RH, regardless of virus solution (Figure 4). These structures became more pronounced, thicker, and fewer in quantity at 50% RH for all virus solutions in pooled human saliva. As expected, crystalline features were not present in virus-laden particles at 80% RH, suggesting particles did not completely effloresce but instead reached an amorphous semi-solid equilibrium state.^60^ Regardless of RH, dried particles always contained some form of a coffee-ring (i.e., accumulation of nonvolatile constituents at the perimeter of a deposited particle^58,59^), though the prominence and thickness of the ring varied by virus type and RH.

**Figure 4.**
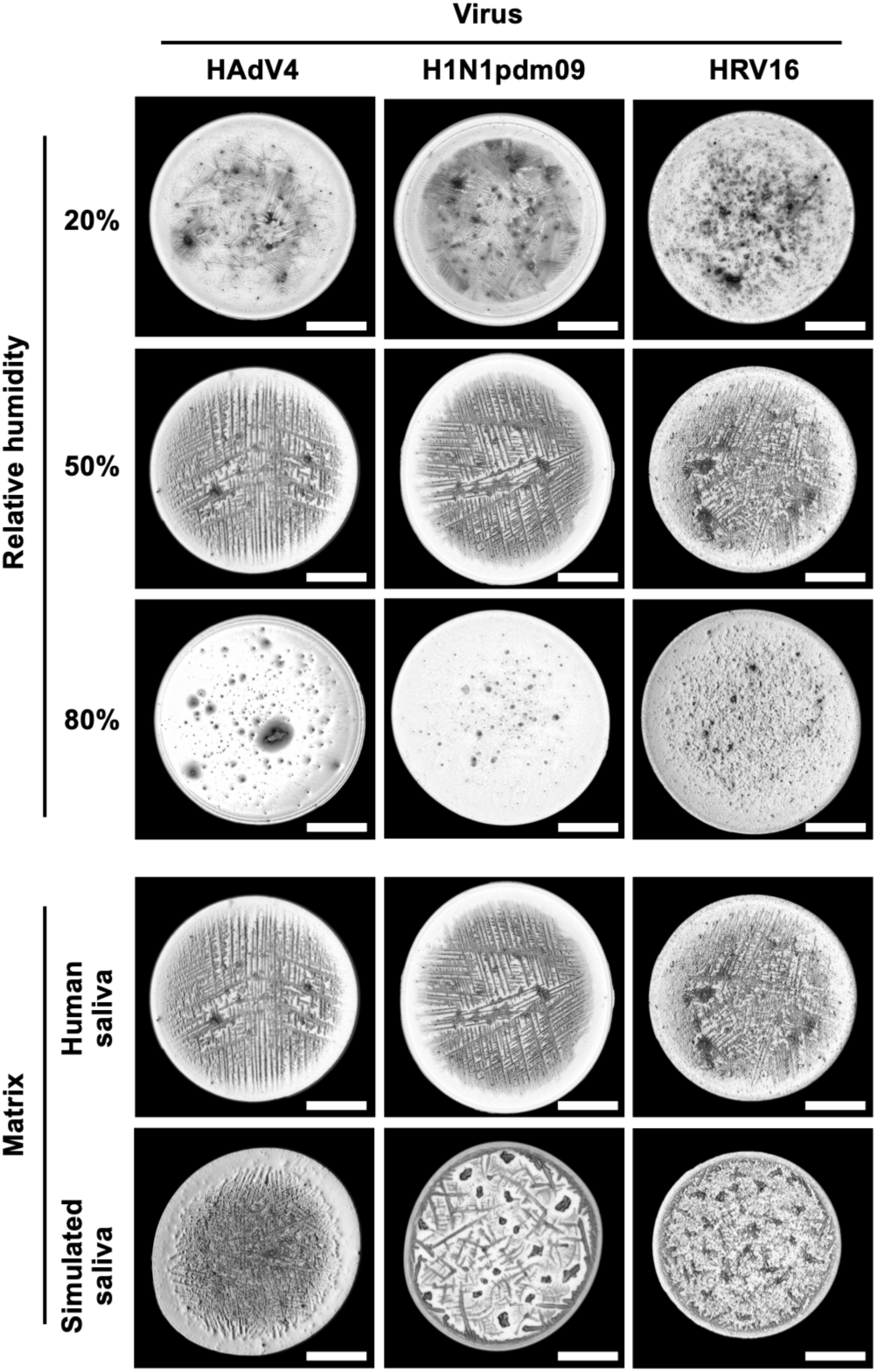
Bright field images of 1 μL deposited pooled human saliva particles containing HAdV4, H1N1pdm09, or HRV16 after drying at 20%, 50%, or 80% RH (top) or bright field images of 1 µL deposited particles comprised of pooled human saliva or simulated saliva containing HAdV4, H1N1pdm09, or HRV16 after drying at 50% RH (bottom). Images were taken at 4x magnification. Scale bars, 500 µm. One representative image is shown for each experimental condition.

Differences in particle composition between simulated and human saliva matrices did not significantly impact drying time of deposited particles, as all particles took approximately 22 min to dry at 50% RH (Table S13). When comparing virus-laden deposited particles of simulated and human saliva at 50% RH, HAdV4-laden particles of simulated and human saliva exhibited relatively comparable morphology with widespread crystalline structures (Figure 4). Interestingly, deposited human saliva particles containing H1N1pdm09 and HRV16 looked markedly different in structure compared to their respective deposited simulated saliva particles after drying (Figure 4). Specifically, simulated saliva particles of H1N1pdm09 showed feathering and far less prominent crystalline structures compared to in human saliva particles. In dried simulated saliva particles of HRV16, a similar phenomenon was observed, where feathering and a lack of elongated crystalline structures were present. These morphologies do not align with the differences observed in decay experiments, indicating that additional factors are driving the extended persistence we see in H1N1pdm09 across human and simulated saliva, in contrast to the distinct trends in HRV16 persistence exhibited in the saliva solutions.

## Discussion

Studies of environmental virus susceptibility in airborne and deposited particles have largely centered on the decay of similarly structured enveloped viruses, including influenza virus and coronavirus, or surrogate viruses, like bacteriophages MS2 and phi6, in laboratory-derived solutions.^3^ Here, we investigated the persistence of three structurally distinct human respiratory viruses in deposited particles of a physiologically relevant matrix, human saliva, at a broad range of RHs commonly experienced in real-world transmission settings. We found that HAdV4, a 100 nm nonenveloped dsDNA virus, exhibited extensive persistence at all RH in deposited human saliva particles, while HRV16, a 30 nm nonenveloped ssRNA virus, was far more susceptible to midrange and elevated RH. H1N1pdm09, a pleomorphic enveloped ssRNA virus, demonstrated greatest persistence at low and mid-range RH, with intermediate degradation at 80% RH compared to the two nonenveloped viruses.

Other work in deposited or airborne suspensions containing influenza virus aligns with the degradation we observed for H1N1pdm09, with reported degradation being minimal at low RH, while decay at mid-range and elevated RH has been described on the order of 0 to 4-log_10_ decay following up to 2 h of exposure in a range of particle matrices.^5,6,14,23–26^ Interestingly, we did not observe the V-shaped viability curve often attributed to influenza virus persistence.^5,9^ This deviation has been noted previously for influenza virus, often in studies using artificial or human respiratory matrices;^6,23,24^ our use of a pooled human saliva matrix could explain the differences we saw here as well. Adenovirus persistence in the limited studies conducted to date has generally been minimal at elevated RH, while observed degradation at low and intermediate RH is greater, ranging from 1- to 4-log_10_ in aerosols or on surfaces with 2.5 h exposure.^10,20^ HAdV4 in our study degraded by at most ∼ 1-log_10_ with 4 h exposure and did not vary with RH. A lack of dependence on midrange and high RH was also observed by Abad et al. over longer timeframes.^21^ Studies have recovered infectious adenovirus off of inoculated indoor surfaces after more than 15 days.^21,61^ Our findings (< 3-log_10_ decay after 24 h) lend further evidence to adenovirus’ persistent nature and indicate the need for mitigation strategies that extend beyond short timeframes.

Over short timeframes (< 2.5 h), previous investigation of rhinovirus decay in aerosolized and deposited particles has established anywhere from negligible to > 2-log_10_ degradation at low, intermediate, and elevated RH.^10,37,62^ Our work falls within this range across RHs up to 2 h post-exposure, but at longer timeframes, our results diverge. HRV16 infectivity dropped below detection (> 4-log_10_ decay) by 6 h at elevated RH in our study, while aerosol and surface persistence of an HRV14 strain was retained for 24 h on surfaces and in aerosols at high RH.^36,62^ Interestingly, Karim and colleagues found that HRV14 infectivity in nasal discharge was lost more rapidly at elevated RH compared to in laboratory derived solutions, including tryptose phosphate broth and a mucin saline suspension,^62^ indicating enhanced rhinovirus decay in respiratory fluids at high RH, as we observed. Of note, rhinovirus is a member of the picornavirus family, which contains several other structurally similar viruses that have been extensively studied. *Picornaviridae* including hepatitis A virus and poliovirus have largely shown increased persistence with elevated RH in aerosols,^11,13,14,16,19^ but trends on surfaces over extended durations show minimal RH effect or increased susceptibility to high RH,^12,22^ similar to what we found for HRV16.

The differential decay of HAdV4 and H1N1pdm09 compared to HRV16 we observed at 50% and 80% RH challenges the commonly held assumption that nonenveloped viruses are more resistant to environmental exposures than enveloped viruses. One potential mechanism driving the faster decay of HRV16 compared to HAdV4 and H1N1pdm09 at intermediate and elevated RH is that HRV16 interacts differently with deposited particle constituents compared to HAdV4 and H1N1pdm09. This could be due to HRV16’s significantly smaller size (i.e., ∼ 3x smaller than H1N1pdm09 and HAdV4) or due to capsid structure and composition, among other differences. Deposited particle morphology was not distinguishable across HAdV4, H1N1pdm09, and HRV16 human saliva suspensions at 50% and 80% RH (Figure 4), confirming that differences in the distribution or behavior of matrix-derived constituents during drying did not occur for distinct virus solutions, as expected. It is possible, however, that virions within these deposited human saliva particles do aggregate or collocate differently, which we could not observe with the brightfield microscopy used in our experiments. While salt constituents inactivate viruses,^13,42,63^ addition of protein constituents can improve virus stability.^5,23^ Differential HRV16 interactions or colocalization with salts or proteins in particles upon reaching equilibrium at 50% or 80% RH could account for the observed differences in inactivation. Further research is needed to investigate molecular interactions of virions with matrix constituents in situ.

Virus decay kinetics in this study were generally well-described by exponential decay curves in one or two phases, with phases split by the drying time of deposited particles at a given RH. Similar decay rates during the drying and equilibrium phases for conditions that resulted in significant virus degradation, like HRV16 at 50% and 80% RH, suggest a mechanism of inactivation driven by virus-constituent interactions that continues during the equilibrium phase. Others have also observed good fit of single phase or biphasic log-linear decay depending on environmental conditions,^6,48^ although these approaches likely oversimplify the complex factors involved in virus degradation with environmental exposure in deposited particles. Continued investigation of mechanistic models^27,44^ to improve model performance is warranted as the field generates additional datasets for a broader range of viruses.

Respiratory matrix composition is an important factor in defining virus persistence. Our pooled human saliva, which has similar properties to previously studied human salivas,^64–66^ provided a physiologically-relevant platform to investigate how viruses persist when expelled in realistic respiratory matrices. Artificial respiratory solutions, like the artificial saliva derived from Woo et al.,^67^ artificial lung fluid, and phosphate buffered saline (PBS) solution, are frequently used as a standardized approach to mimic respiratory solutions. Importantly, these laboratory-derived matrices can have elevated levels of salt and proteins compared to real-world respiratory solutions. PBS, for example, typically contains ∼10 g/L of salt, roughly an order of magnitude more salt than we measured in our saliva. Artificial saliva solutions exhibit upwards of 5x more protein than we detected in our saliva, with ∼3 g/L proteins commonly added.^60,67,68^ Artificial lung fluid often consists of an even higher amount of protein than in artificial saliva,^23,69–71^ with protein levels stemming from estimates extrapolated from bronchial alveolar lavage (BAL) fluids. More work is needed to understand the relative contribution of different respiratory matrices that end up in the emitted particles contributing to onward transmission, as this would inform the solutions that researchers should use to most accurately study environmental virus persistence.

Here, we used the same quantities of protein and salt measured in the pooled human saliva, 0.73 g/L protein, and 2.2 g/L salt, to generate a protein-salt matrix that recapitulated the composition of saliva. Using this solution, we were able to reproduce the stability of HAdV4 and H1N1pdm09 at 50% RH in human saliva, but we observed significantly greater HRV16 persistence in the simulated saliva matrix than in human saliva at 4 h. Differential effects of matrix composition on influenza virus stability at variable organic-to-salt ratios have previously been observed, with albumin:NaCl ratios above 0.11:1 conferring more protection.^23^ Our organic:salt ratio, 0.33:1, although above the threshold defined for protection of influenza virus, was the same in both simulated and human matrices, and this factor was therefore not governing the differential trends we observed in HRV16 persistence.

Importantly, protein type, not just concentration, plays a role in driving virus protection. For example, Schaub et al. found that smaller proteins, including chicken albumin, stabilized influenza virus more effectively than larger proteins, like ferritin, at the same mass concentration.^23^ For the protein in our simulated saliva, we used human serum albumin, an intermediate-sized protein (∼ 69 kDa^57^) prevalent in human saliva.^72^ It is possible that the protective protein effect is more pronounced for virions of different sizes, and the much smaller size of HRV16 compared to that of H1N1pdm09 or HAdV4 could alter interactions with proteins of a particular size range. In our pooled human saliva, which comprised a range of proteins, including high abundance of large proteins like mucin 5B (∼596 kDa^57^), we propose that larger proteins were effective in protecting H1N1pdm09 and HAdV4 but not HRV16, and once all protein content was comprised of smaller protein in the simulated saliva, HRV16 was also protected. Additional work is needed using a range of simulated salivas containing human proteins of variable sizes to evaluate this hypothesis.

Salt and protein content can also impact the drying time of deposited particles, and time to efflorescence contributes to virus persistence.^23^ In our study, we did not see any significant differences in the drying times of our simulated or human salivas across the viruses used. Interestingly, we did observe an increased ‘coffee ring’ region in the simulated saliva particles of H1N1pdm09 and HRV16 compared to those of human saliva, as well as fewer crystalline lattice structures throughout the dried particle at 50% RH. Addition of proteins like albumin to particles has reduced the formation of large crystals and enhanced constituent mixing, resulting in a more homogeneous particle morphology with fewer crystalline structures.^68^ Our simulated saliva particles appeared to show this morphology, while the human saliva did not. This suggests the presence or absence of proteins or other organic constituents in our pooled saliva that we were not able to recapitulate with the simple formula we used for our simulated saliva.

Our study has limitations. First, we were unable to fully recapitulate the phenotypes observed in real human saliva with our simplified, simulated saliva. Importantly, we used a single human protein and a mixture of salts in our simulated saliva, and the inclusion of more constituents, including lipids, other organics, and additional proteins that better correspond to those found in human saliva would allow for a simulated saliva that better mimics the trends observed for HRV16 decay in human saliva. Balancing the simplicity of a standardized solution with the complexity of real-world respiratory matrices will be critical for developing representative but feasible solutions that represent viral emissions and allow for accurate determination of environmental virus persistence. Second, variability in saliva composition exists, and here, although we studied a pooled sample to encompass differences across individuals, it is likely that other salivas have discrete characteristics that could differentially impact virus persistence. Indeed, our pooled human saliva matrix was more protective of H1N1pdm09, by about 1.5-log_10_, from previous work done in the same experimental model using a different pooled saliva sample.^6^ Respiratory fluid composition can vary during an infection,^73,74^ and use of saliva from sick individuals could provide further insights beyond our use of saliva from healthy participants. Another limitation of our work is the use of 1 µL deposited particles. These are on the large end of particle sizes emitted during human respiratory activities;^40^ particles in the smaller size ranges merit further investigation, as particle size can result in differences in virus stability outside the host,^48^ and particles of other sizes are likely also important in contributing to onward transmission.^1,2^

### Environmental implications

In this study, we showed that distinct human viruses contained in deposited human saliva particles respond differently to environmental exposures, highlighting that mitigation strategies cannot rely on a single-virus approach to effectively reduce infection risks. Crowded indoor settings, such as schools or public transit, for example, may benefit from optimized RH management that can reduce infectious virus loads during seasonal outbreaks of HRV16 or H1N1pdm09, while RH-based approaches may not be as useful for reducing surface-associated HAdV4 burden. Our findings also underscore the need to move beyond overgeneralizations of trends in enveloped versus nonenveloped virus environmental susceptibility. The species-specific nature of environmental virus degradation demands the in-depth study of a broad range of human viruses, as multiple virus-related factors, such as viral protein composition, host cell entry mechanisms, or surface antigens, could contribute to persistence. In addition, we emphasized the need for a simulated respiratory matrix that will accurately recapitulate virus degradation in emitted respiratory solutions. Future work should explore a range of physiologically relevant or representative matrices and work to define the underlying mechanisms governing virus stability during the environmental phase of respiratory virus spread. Overall, our findings contribute to foundational environmental persistence data that are critical in developing comprehensive public health guidance.

## Data availability

All data will be made available on the Duke Research Data Repository upon publication.

## Supporting Information Available

The Supporting information is available free of charge.

Supporting information text includes details of virus and mammalian cell culture, proteomics sample preparation, processing, and data analysis, and details of extractions and (RT)-qPCR conducted for nucleic acid recovery determination; tables include virus characteristics, pooled human saliva donor information, infectious virus stock concentrations, (RT-)qPCR assay details, statistical analyses, model performance, regression comparisons, physicochemical properties of saliva and simulated saliva, virus stock composition, and infectious virus decay data in bulk controls, and protein group abundance data; figures include temperature and relative humidity profiles during deposited particle experiments, infectious virus decay data, genome copy loss, deposited particle drying times, and model fit curves.

## Acknowledgements

This work was supported in part by the Engineering Research Centers Program of the National Science Foundation under NSF Cooperative Agreement No. EEC-2133504, the Swiss National Science Foundation (grant P500PN_230569), and FluLab. Proteomics were conducted through the Duke Proteomics and Metabolomics Core Facility by Dr. Matthew Foster. ICP-MS was conducted in the Hsu-Kim laboratory by Dr. Nelson Rivera.

The authors thank members of the Rockey Research Group for careful discussions and feedback in improving the text.

