## Supporting Information for "Relative humidity differentially modulates decay of structurally distinct respiratory viruses in deposited human saliva particles"

This PDF file includes:

Supporting information text
Tables S1 to S13
Figures S1 to S6

Total pages: 20
Total number of tables: 13
Total number of figures: 6

### Supporting information text

***Virus propagation and stock preparation.*** HAdV4 propagation was conducted by growing virus at a multiplicity of infection (MOI) = 0.1 on 90% A549 cells (ATCC, Cat. No. CCL-185) in infection medium (DMEM containing high glucose and supplemented with 1% L-glutamine and 1% penicillin-streptomycin) at 37°C and 5% $CO_2$. HAdV4 was harvested after significant cytopathic effect was observed at approximately 6 days post-inoculation (dpi) by conducting three rounds of freeze–thawing, followed by removal of cellular debris through centrifugation at 2,500 x g for 10 minutes at 4°C.

H1N1pdm09 propagation was conducted by growing virus at MOI = 0.001 on confluent Madin-Darby Canine Kidney (MDCK) cells, kindly provided by Dr. Seema Lakdawala, in infection medium (MEM supplemented with 2% antibiotic-antimycotic (Thermo Fisher Scientific, Cat. No. 15240062), 1% L-glutamine, and 1 µg/mL TPCK trypsin (Worthington-Biochemicals, Cat. No. LS003750)) at 37°C and 5% $CO_2$. H1N1pdm09 was harvested at 24 hours post-inoculation (hpi), and cellular material was removed through centrifugation at 2,500 x g for 10 minutes at 4°C.

HRV16 was propagated at MOI = 1 in 90% confluent H1HeLa cells (ATCC, CRL-1958) in infection medium (DMEM with high glucose, supplemented with 1% L-glutamine and 1% penicillin-streptomycin) at 33°C and 5% $CO_2$. HRV16 was harvested following significant cytopathic effect, at approximately 3 - 4 dpi, by scraping cells and centrifuging at 2,500 × g for 10 minutes at 4 °C. All but 5 mL of the supernatant was collected and kept on ice, leaving a small portion of supernatant above the cell pellet. The pellet underwent three rounds of freeze–thawing and was subsequently centrifuged at 2,500 x g for 10 minutes at 4°C, and the resulting supernatant was pooled with the initial fraction.

Following virus propagation and harvesting, all viruses were concentrated through a 0.22 µm-filtered 30% sucrose cushion (Sigma-Aldrich, Cat. No. S0389) by ultracentrifugation at 24,000 rpm for 2.5 hours at 4°C on a SW 32 Ti rotor in a Beckman Coulter Optima XPN-80 ultracentrifuge. Supernatant was removed, and residual HAdV4 and HRV16 pellets were washed once with PBS or the corresponding matrix before resuspension; H1N1pdm09 pellets were not washed. Two solutions were used for virus stock resuspension, namely pooled human saliva (Cell Sciences, Cat. No. CSI20375B) and a simulated saliva matrix. Purchased pooled human saliva consisted of saliva from 10 individuals (Table S2). No pre-treatment of the pooled human saliva was carried out by the supplier or the laboratory prior to use for virus resuspension. Simulated saliva was created to recapitulate the protein and salt concentrations characterized in the pooled human saliva. Briefly, simulated saliva was comprised of 0.547 mg x $mL^{-1}$ NaCl, 1.606 mg x $mL^{-1}$ KCl, 0.023 mg x $mL^{-1}$ $CaCl_2$, 0.009 mg x $mL^{-1}$ $MgCl_2$, and 730 µg/mL human serum albumin (Albumin, Human Serum, Fraction V, High Purity, Millipore Sigma, Cat. No. 126658) dissolved in autoclaved purified water (i.e., milliQ water). Virus stocks were aliquoted and stored at -80°C until use. Infectious virus concentrations were quantified using the 50% tissue culture infectious dose ($TCID_{50}$) Spearman–Kärber assay as previously described.[1] The concentration of virus stocks in human and simulated saliva used in experiments ranged from ~ 2 x $10^7$ $TCID_{50}$/mL to 6 x $10^7$ $TCID_{50}$/mL (Table S3).

***Human saliva protein profile.*** To prepare the pooled human saliva for proteomics analysis, 10 µg of saliva was adjusted to 5% SDS (w/v) and 50 mM triethylammonium bicarbonate, pH 8.5 (TEAB) followed by addition of 10 mM dithiothreitol and heating at 80 °C for 10 min on a Thermomixer. After cooling, 25 mM iodoacetamide was added followed by incubation in the dark at room temp for 30 min. Detergent removal and trypsin digestion were performing using an S-trap micro device (Protifi) using manufacturer's instructions. The trypsin digestion used 1 µg of Sequencing grade modified trypsin (Promega) at 47 °C for 1 h. After peptide elution and lyophilization, peptides were reconstituted in 0.1% formic acid, and 500 ng of digest was loaded onto a disposable trap column (Evotip Pure). Liquid chromatograph coupled to tandem mass spectrometry (LC-MS/MS) was performed using an Evosep One LC interfaced to a ThermoFisher Orbitrap Astral. The Evosep used a 30 sample-per-day method with a Bruker Pepsep 15 cm x 150 µm column (1.5 µm particle size) and a PepSep Sprayer and stainless steel (30 µm) emitter. MS/MS used a 240,000-resolution precursor scan m/z 380-980, an AGC target of 500% and 0.6 s cycle time. Astral MS/MS used fixed windows of 4 m/z from 380-80 m/z, target AGC of 500%, max fill time of 6 ms and a normalized collision energy of 28%.

Raw MS data was converted to *.htrms format using HTRMS converter and processed in Spectronaut 20.1 (Biognosys). A spectral library generated by direct-DIA search of a Homo sapiens consensus database[2] and appended with contaminant sequences using FragPipe, as well as a 3 bacterium database as described previously.[3] Default settings were used except no variable modifications were selected. The resulting library was used to analyze the data in Spectronaut at a 1% precursor and 1% protein group false discovery rate.

***Nucleic acid extraction and (RT)-qPCR.*** Nucleic acid extractions were conducted with 140 µL of each sample using the QIAmp Viral RNA Mini Kit (Qiagen, Cat. No. 52906) following the manufacturer's instructions. Carrier RNA was included in extractions of the RNA viruses, H1N1pdm09 and HRV16. Nucleic acids were eluted in 60 µL AVE buffer and stored at -80°C until further analysis. Negative extraction controls, comprising 140 µL nuclease-free water were included in each extraction. Reverse transcription quantitative polymerase chain reaction (RT-qPCR) and quantitative polymerase chain reaction (qPCR) analyses were performed using the iTaq Universal Probes One-Step Kit (BioRad, Cat. No. 1725141) and the iTaq Universal Probes Supermix (BioRad, Cat. No. 1725131), respectively. Thermocycling was performed on a CFX Opus 96 real-time PCR system (BioRad), and Cq values were determined using CFX Maestro software (BioRad). All assays were conducted in 20 µL reaction volumes, with primer and probe concentrations of 0.5 µM and 0.15 µM, respectively. 5 µL of template were included in each reaction. RT-qPCR assays included an initial reverse transcription step at 50°C for 10 min, followed by initial denaturation at 95°C for 1 min, while the qPCR assay included an initial denaturation at 95°C for 1 min. Following initial denaturation, both RT-qPCR and qPCR cycling conditions included 40 cycles of denaturation at 95°C for 10 s, followed by annealing/extension at 60°C for 20 s.

In vitro transcribed RNA was generated for HRV16 and H1N1pdm09 standards, and purified PCR product was used for the HAdV4 standard. All samples, standards, and no template controls were run in duplicate, and standard curves comprised six ten-fold serial dilutions of the highest standard. For H1N1pdm09 and HRV16 standards, in vitro transcribed RNA of the full-length A/CA/04/2009 (H1N1) M gene (Accession No. MN371612) and the full-length RVA1a 5′ untranslated region (5′ UTR; Accession No. NC_03831) were used, respectively. In vitro transcription was carried out using the MEGAscript T7 Transcription kit (Thermofisher, Cat. No. AM1333). For the HAdV4 standard, a purified PCR product comprising the full length Hexon gene (Accession No. KX384949.1) was used. Additional details of RT-qPCR and qPCR assays are included in Table S4.

### Supporting information tables

**Table S1.** Properties of the human respiratory viruses used in this study.

| Virus | Family | Size | Morphology | Genome |
|---|---|---|---|---|
| **Adenovirus type 4 (HAdV4)** | *Adenoviridae* | ~90 nm | non-enveloped icosahedral | dsDNA, ~36 kbp, linear |
| **Influenza A virus A/CA/07/2009 (H1N1) (H1N1pdm09)** | *Orthomyxoviridae* | ~100 to 300 nm | enveloped spherical to filamentous | (-) ssRNA, ~13.5 kb, linear, segmented |
| **Rhinovirus A type 16 (HRV16)** | *Picornaviridae* | ~30 nm | non-enveloped icosahedral | (+) ssRNA, ~7.1 kb, linear |

nm = nanometers, kb = kilobases, kbp = kilobase pairs, ssRNA = single-stranded RNA, dsDNA = double-stranded DNA.

**Table S2.** Donor information for the pooled human saliva purchased from Cell Sciences (Catalog No. CSI20375B).

| Donor | Birth Year | Gender | Race |
|---|---|---|---|
| T1866 | 1968 | Male | Caucasian |
| T1995 | 1986 | Male | Caucasian |
| T5234 | 1937 | Male | Caucasian |
| T5717 | 1978 | Female | Caucasian |
| T5729 | 1979 | Male | Caucasian |
| T6445 | 1962 | Male | Caucasian |
| T6551 | 1960 | Female | Caucasian |
| T6735 | 1998 | Female | African American/Caucasian |
| T6736 | 2000 | Male | Caucasian |
| T6810 | 1985 | Female | Caucasian |

**Table S3.** Infectious virus concentrations, in $TCID_{50}$/mL, in pooled human saliva and simulated saliva stocks.

| **HAdV4** | | **H1N1pdm09** | | **HRV16** | |
|---|---|---|---|---|---|
| Human saliva | Simulated saliva | Human saliva | Simulated saliva | Human saliva | Simulated saliva |
| $6.1 \times 10^7$ | $8.9 \times 10^8$ | $3.8 \times 10^7$ | $5.0 \times 10^7$ | $1.6 \times 10^7$ | $1.2 \times 10^7$ |

**Table S4.** RT-qPCR and qPCR assay details, including primers and probe sequences, target amplicon size, reaction volume and cycling conditions, and assay slope, efficiency, and goodness of fit ($R^2$).

| Virus | Primer sequence (5´ to 3´) | Amplicon size (bp) | Probe sequence | Standard curve | | |
|---|---|---|---|---|---|---|
| | | | | Slope | Efficiency (%) | $R^2$ |
| HAdV4[4] | F: CKT ACA TGC ACA TCK CSG G<br>R: GCA TTY TGG ACA AAV CGT CTA CG | 68 | 5HEX / CCG GRC TCA / ZEN / GGT ACT CCG ARG CGT CCT / 3IABkFQ | -3.28 | 105 | 0.99 |
| H1N1pdm09[5] | F: CAA GAC CAA TCY TGT CAC CTC TGA C<br>R: GCA TTY TGG ACA AAV CGT CTA CG | 106 | 56-FAM / TGC AGT CCT / ZEN / CGC TCA CTG GGC ACG / 3IABkFQ | -3.90 | 80.63 | 0.999 |
| HRV16[6] | F: GTG TGA AGA GCC GCG TGT<br>R: TGG CTG CAG GTT TAA GGT TAGC | 72 | 5ATTO590N / TCC TCC GGC CCC TGA ATG YGG C / 3IAbRQSp | -3.82 | 82.9 | 0.998 |

HEX = hexachlorofluorescein; ZEN = ZEN quencher; 3IABkFQ = 3' Iowa Black FQ dark quencher; 3IAbRQSp = 3' Iowa Black RQ dark quencher

**Table S5.** One-way ANOVA with post hoc multiple comparisons of virus decay in saliva across relative humidities for each virus and exposure time.

| Virus | Exposure time | Tukey's multiple comparisons test | Mean difference | 95% confidence interval of difference | Adjusted p-value |
|---|---|---|---|---|---|
| **HAdV4** | 1h | 20% vs 50% | -0.1973 | -0.5794 to 0.1849 | 0.3619 |
| | | 20% vs 80% | -0.4596 | -0.8947 to -0.02447 | 0.0391 |
| | | 50% vs 80% | -0.2623 | -0.6784 to 0.1537 | 0.2366 |
| | 2h | 20% vs 50% | 0.1673 | -0.3477 to 0.6822 | 0.6498 |
| | | 20% vs 80% | -0.5338 | -1.120 to 0.05257 | 0.0738 |
| | | 50% vs 80% | -0.701 | -1.262 to -0.1404 | 0.0169 |
| | 4h | 20% vs 50% | 0.0475 | -0.4645 to 0.5595 | 0.9638 |
| | | 20% vs 80% | -0.0025 | -0.5854 to 0.5804 | >0.9999 |
| | | 50% vs 80% | -0.05 | -0.6074 to 0.5074 | 0.9661 |
| | 6h | 20% vs 50% | 0.681 | -0.3971 to 1.759 | 0.2355 |
| | | 20% vs 80% | 0.4917 | -0.7358 to 1.719 | 0.5273 |
| | | 50% vs 80% | -0.1893 | -1.363 to 0.9843 | 0.8955 |
| **H1N1pdm09** | 1h | 20% vs 50% | 0.5208 | -0.3039 to 1.346 | 0.2198 |
| | | 20% vs 80% | 0.335 | -0.5467 to 1.217 | 0.5334 |
| | | 50% vs 80% | -0.1858 | -1.011 to 0.6389 | 0.791 |
| | 2h | 20% vs 50% | 0.76 | 0.04343 to 1.477 | 0.0393 |
| | | 20% vs 80% | 0.9167 | 0.1506 to 1.683 | 0.0231 |
| | | 50% vs 80% | 0.1567 | -0.5599 to 0.8732 | 0.8016 |
| | 4h | 20% vs 50% | 0.4967 | -0.3742 to 1.368 | 0.2635 |
| | | 20% vs 80% | 0.915 | 0.04413 to 1.786 | 0.0413 |
| | | 50% vs 80% | 0.4183 | -0.4525 to 1.289 | 0.3664 |
| | 6h | 20% vs 50% | 1.165 | -0.06534 to 2.395 | 0.0612 |
| | | 20% vs 80% | 2.293 | 1.063 to 3.524 | 0.003 |
| | | 50% vs 80% | 1.128 | -0.1020 to 2.359 | 0.0686 |
| **HRV16** | 1h | 20% vs 50% | -1.208 | -2.144 to -0.2728 | 0.0174 |
| | | 20% vs 80% | -1.252 | -2.187 to -0.3161 | 0.0149 |
| | | 50% vs 80% | -0.04333 | -0.9789 to 0.8922 | 0.9889 |
| | 2h | 20% vs 50% | -1 | -2.127 to 0.1268 | 0.0769 |
| | | 20% vs 80% | -0.96 | -2.087 to 0.1668 | 0.0883 |
| | | 50% vs 80% | 0.04 | -1.087 to 1.167 | 0.9935 |
| | 4h | 20% vs 50% | 0.9583 | -0.08907 to 2.006 | 0.0692 |
| | | 20% vs 80% | 1.253 | 0.2059 to 2.301 | 0.0243 |
| | | 50% vs 80% | 0.295 | -0.7524 to 1.342 | 0.6803 |
| | 6h | 20% vs 50% | 2.042 | 0.9336 to 3.150 | 0.0032 |
| | | 20% vs 80% | 2.498 | 1.390 to 3.606 | 0.0011 |
| | | 50% vs 80% | 0.4567 | -0.6514 to 1.565 | 0.4627 |

**Table S6.** One-way ANOVA with post hoc multiple comparisons of virus decay in saliva across viruses for each relative humidity and exposure time.

| Relative humidity | Exposure time | Tukey's multiple comparisons test | Mean difference | 95% confidence interval of difference | Adjusted p-value |
|---|---|---|---|---|---|
| 20% | 1h | HAdV4 vs. H1N1pdm09 | -0.3346 | -1.236 to 0.5665 | 0.5474 |
| | | HAdV4 vs. HRV16 | 0.9571 | 0.05599 to 1.858 | 0.039 |
| | | H1N1pdm09 vs. HRV16 | 1.292 | 0.3284 to 2.255 | 0.0134 |
| | 2h | HAdV4 vs. H1N1pdm09 | -0.8238 | -1.616 to -0.03119 | 0.0427 |
| | | HAdV4 vs. HRV16 | 0.8863 | 0.09369 to 1.679 | 0.0313 |
| | | H1N1pdm09 vs. HRV16 | 1.71 | 0.8627 to 2.557 | 0.0014 |
| | 4h | HAdV4 vs. H1N1pdm09 | -0.2492 | -1.189 to 0.6905 | 0.7257 |
| | | HAdV4 vs. HRV16 | 0.7892 | -0.1505 to 1.729 | 0.0956 |
| | | H1N1pdm09 vs. HRV16 | 1.038 | 0.03376 to 2.043 | 0.0437 |
| | 6h | HAdV4 vs. H1N1pdm09 | -0.26 | -1.279 to 0.7595 | 0.7426 |
| | | HAdV4 vs. HRV16 | 1.032 | 0.01221 to 2.051 | 0.0477 |
| | | H1N1pdm09 vs. HRV16 | 1.292 | 0.2018 to 2.382 | 0.0241 |
| 50% | 1h | HAdV4 vs. H1N1pdm09 | 0.3835 | -0.1585 to 0.9255 | 0.1739 |
| | | HAdV4 vs. HRV16 | -0.054 | -0.6440 to 0.5360 | 0.9648 |
| | | H1N1pdm09 vs. HRV16 | -0.4375 | -1.055 to 0.1796 | 0.1729 |
| | 2h | HAdV4 vs. H1N1pdm09 | -0.231 | -0.8583 to 0.3963 | 0.5788 |
| | | HAdV4 vs. HRV16 | -0.281 | -0.9639 to 0.4019 | 0.5104 |
| | | H1N1pdm09 vs. HRV16 | -0.05 | -0.7642 to 0.6642 | 0.9792 |
| | 4h | HAdV4 vs. H1N1pdm09 | 0.2 | -0.3472 to 0.7472 | 0.5718 |
| | | HAdV4 vs. HRV16 | 1.7 | 1.153 to 2.247 | <0.0001 |
| | | H1N1pdm09 vs. HRV16 | 1.5 | 0.8883 to 2.112 | 0.0003 |
| | 6h | HAdV4 vs. H1N1pdm09 | 0.224 | -1.020 to 1.468 | 0.8666 |
| | | HAdV4 vs. HRV16 | 2.392 | 1.148 to 3.636 | 0.0015 |
| | | H1N1pdm09 vs. HRV16 | 2.168 | 0.7774 to 3.559 | 0.0053 |
| 80% | 1h | HAdV4 vs. H1N1pdm09 | 0.46 | -0.07678 to 0.9968 | 0.0866 |
| | | HAdV4 vs. HRV16 | 0.165 | -0.3718 to 0.7018 | 0.6355 |
| | | H1N1pdm09 vs. HRV16 | -0.295 | -0.8318 to 0.2418 | 0.2848 |
| | 2h | HAdV4 vs. H1N1pdm09 | 0.6267 | -0.2447 to 1.498 | 0.1484 |
| | | HAdV4 vs. HRV16 | 0.46 | -0.4114 to 1.331 | 0.3086 |
| | | H1N1pdm09 vs. HRV16 | -0.1667 | -1.038 to 0.7047 | 0.832 |
| | 4h | HAdV4 vs. H1N1pdm09 | 0.6683 | -0.1717 to 1.508 | 0.11 |
| | | HAdV4 vs. HRV16 | 2.045 | 1.205 to 2.885 | 0.0007 |
| | | H1N1pdm09 vs. HRV16 | 1.377 | 0.5366 to 2.217 | 0.0057 |
| | 6h | HAdV4 vs. H1N1pdm09 | 1.542 | 0.3660 to 2.717 | 0.0163 |
| | | HAdV4 vs. HRV16 | 3.038 | 1.863 to 4.214 | 0.0005 |
| | | H1N1pdm09 vs. HRV16 | 1.497 | 0.3210 to 2.672 | 0.0186 |

**Table S7.** Drying times of deposited human saliva particles containing HAdV4, H1N1pdm09 or HRV16 exposed to 20%, 50%, or 80% relative humidity (RH). Drying times were determined by taking the mean drying time of the first and last deposited 10 x 1 µL particles to dry. Two independent drying experiments were conducted per condition.

| **RH** | **Virus** | **Drying time, min** Mean (minimum to maximum) |
|---|---|---|
| 20% | HAdV4 | 16.39 (16.38 to 16.40) |
| | H1N1pdm09 | 15.03 (14.52 to 15.54) |
| | HRV16 | 14.09 (14.02 to 14.16) |
| 50% | HAdV4 | 23.38 (21.55 to 25.21) |
| | H1N1pdm09 | 20.36 (16.40 to 24.31) |
| | HRV16 | 20.36 (16.14 to 24.58) |
| 80% | HAdV4 | 48.99 (47.41 to 50.56) |
| | H1N1pdm09 | 50.29 (50.2 to 50.38) |
| | HRV16 | 41.82 (41.29 to 42.34) |

**Table S8.** Performance indicators of two different models describing HAdV4, H1N1pdm09, and HRV16 decay at each relative humidity (RH). The two models tested are a single-phase linear model and a two-phase linear model that uses deposited particle drying time as the breakpoint.

| Virus | RH | Single-phase linear model | | Two-phase linear model | |
|---|---|---|---|---|---|
| | | **Performance indicator** | **Value** | **Performance indicator** | **Value** |
| **HAdV4** | 20% | $R^2$ (Goodness of fit) | 0.32 | $R^2$ | 0.54 |
| | | Slope (Decay rate) | -0.002 | Slope wet | -0.032 |
| | | 95% CI | -0.002497 to -0.0006616 | 95% CI | -0.05072 to -0.01406 |
| | | | | Slope dry | -0.001 |
| | | | | 95% CI | -0.001830 to -0.0001334 |
| | 50% | $R^2$ | 0.62 | | 0.63 |
| | | Slope | -0.004 | Slope wet | -0.014 |
| | | 95% CI | -0.004654 to -0.002622 | 95% CI | -0.03139 to 0.002946 |
| | | | | Slope dry | -0.003 |
| | | | | 95% CI | -0.004442 to -0.002108 |
| | 80% | $R^2$ | 0.80 | $R^2$ | 0.82 |
| | | Slope | -0.003 | Slope wet | 0.000552 |
| | | 95% CI | -0.004079 to -0.002493 | 95% CI | -0.005870 to 0.006974 |
| | | | | Slope dry | -0.003714 |
| | | | | 95% CI | -0.004771 to -0.002656 |
| **H1N1pdm09** | 20% | $R^2$ | 0.28 | $R^2$ | 0.28 |
| | | Slope | -0.002 | Slope wet | 0.000 |
| | | 95% CI | -0.002900 to -0.0003647 | 95% CI | -0.03310 to 0.03313 |
| | | | | Slope dry | -0.002 |
| | | | | 95% CI | -0.003076 to -0.0002428 |
| | 50% | $R^2$ | 0.7138 | $R^2$ | 0.7392 |
| | | Slope | -0.004 | Slope wet | -0.019 |
| | | 95% CI | -0.005460 to -0.003071 | 95% CI | -0.03934 to 0.002253 |
| | | | | Slope dry | -0.004 |
| | | | | 95% CI | -0.005155 to -0.002471 |
| | 80% | $R^2$ | 0.91 | $R^2$ | 0.9112 |
| | | Slope | -0.007 | Slope wet | -0.006234 |
| | | 95% CI | -0.008388 to -0.006198 | 95% CI | -0.01520 to 0.002734 |

| | | | | | |
|---|---|---|---|---|---|
| **H1N1pdm09** | 80% | | | Slope dry | -0.007415 |
| | | | | 95% CI | -0.008944 to -0.005887 |
| **HRV16** | 20% | $R^2$ | 0.27 | $R^2$ | 0.6437 |
| | | Slope | -0.003 | Slope wet | -0.092 |
| | | 95% CI | -0.004990 to -0.0007279 | 95% CI | -0.1343 to -0.04879 |
| | | | | Slope dry | -0.001 |
| | | | | 95% CI | -0.003100 to 0.0003255 |
| | 50% | $R^2$ | 0.95 | $R^2$ | 0.9595 |
| | | Slope | -0.011 | Slope wet | 0.005333 |
| | | 95% CI | -0.01225 to -0.009911 | 95% CI | -0.01685 to 0.02752 |
| | | | | Slope dry | -0.01154 |
| | | | | 95% CI | -0.01282 to -0.01025 |
| | 80% | $R^2$ | 0.93 | $GR^2$ | 0.95 |
| | | Slope | -0.012 | Slope wet | 0.004 |
| | | 95% CI | -0.01404 to -0.01079 | 95% CI | -0.009632 to 0.01833 |
| | | | | Slope dry | -0.014 |
| | | | | 95% CI | -0.01575 to -0.01199 |

**Table S9.** F-test results for comparisons of linear model slopes.

| | **F** | **DFn** | **DFd** | **p-value** |
|---|---|---|---|---|
| *Comparison of linear slopes per virus among different RH* | | | | |
| HAdV4 50% vs 80% | 0.245 | 1 | 52 | 0.6227 |
| H1N1pdm09 20% vs 50% | 9.957 | 1 | 41 | 0.003 |
| H1N1pdm09 20% vs 80% | 50.03 | 1 | 38 | <0.0001 |
| H1N1pdm09 50% vs 80% | 14.97 | 1 | 41 | 0.0004 |
| HRV16 50% vs 80% | 1.957 | 1 | 38 | 0.17 |
| *Comparison of linear slopes per RH among different viruses* | | | | |
| HAdV4 vs H1N1pdm09 50% | 0.6527 | 1 | 55 | 0.4226 |
| HAdV4 vs HRV16 50% | 91.71 | 1 | 52 | 0.0001 |
| H1N1pdm09 vs HRV16 50% | 71.84 | 1 | 41 | 0.0001 |
| AdV4 vs HRV16 80% | 112 | 1 | 38 | P<0.0001 |
| AdV4 vs H1N1pdm09 80% | 38.49 | 1 | 38 | P<0.0001 |
| *Is the slope statistically significantly different from 0?* | | | | |
| HAdV4 20% dry phase versus 0 | 5.68 | 1 | 25 | 0.0251 |
| HRV16 20% dry phase versus 0 | 2.896 | 1 | 18 | 0.1060 |

**Table S10.** Protein groups and their relative abundance determined via proteomics in the pooled human saliva.

*This table is provided as an external data file.*

**Table S11.** Total protein concentration, total sucrose concentration, and elemental concentrations of virus stocks in human saliva. Two independent measurements were conducted for each analysis.

| | **HAdV4** mean (min to max) | **H1N1pdm09** mean (min to max) | **HRV16** mean (min to max) |
|---|---|---|---|
| **Total protein concentration, μg/mL** | 846.65 (838.31 to 854.98) | 830.96 (824.19 to 837.73) | 874.43 (860.53 to 888.32) |
| **Total sucrose concentration, mg/mL** | 0.57 (0.56 to 0.58) | 1.65 (1.63 to 1.66) | 1.22 (1.22 to 1.22) |
| **Elemental concentration, mg/L** | | | |
| **Na** | 244.18 (238.75 to 249.61) | 238.85 (232.87 to 244.82) | 241.60 (238.35 to 244.84) |
| **Mg** | 2.47 (2.42 to 2.51) | 2.39 (2.36 to 2.42) | 2.60 (2.59 to 2.60) |
| **K** | 842.02 (824.26 to 859.77) | 835.95 (813.39 to 858.51) | 865.59 (858.92 to 872.25) |
| **Ca** | 9.85 (9.21 to 10.49) | 9.10 (8.70 to 9.49) | 11.34 (10.04 to 12.63) |

**Table S12.** Statistical comparison of infectious virus loss in deposited particles of human saliva compared to simulated saliva after different exposure times. unpaired t-tests under the assumption of homogeneity in variance (Figure 3). The significance of differences among variances were tested using a F test and Welch's correction was applied accordingly.

| **Virus** | **Exposure time, hours** | **t(df) = t-value** | **p-value** |
|---|---|---|---|
| HAdV4 | 1 | t(6) = 2.006 | 0.0916 |
| | 2 | t(6) = 0.7526 | 0.4802 |
| | 4 | t(6) = 0.7023 | 0.5088 |
| H1N1pdm09 | 1 | t(6) = 0.9869 | 0.3690 |
| | 2 | t(5) = 1.344 | 0.2366 |
| | 4 | t(4) = 1.942 | 0.1240 |
| HRV16 | 1 | t(2.001) = 2.615 | 0.1203 |
| | 2 | t(1.4) =0.3050 | 0.7995 |
| | 4 | t(2.985) = 9.859 | 0.0023 |

**Table S13.** Drying times of deposited human saliva and simulated saliva particles containing HAdV4, H1N1pdm09, or HRV16 at 50% RH. Drying times were determined by taking the mean drying time of the first and last deposited 10 x 1 µL particles to dry. Two independent drying experiments were conducted per condition.

| **Matrix** | **Virus** | **Drying time, min**<br>Mean (minimum to maximum) |
|---|---|---|
| Human saliva | HAdV4 | 23.38 (21.55 to 25.21) |
| | H1N1pdm09 | 20.36 (16.40 to 24.31) |
| | HRV16 | 20.36 (16.14 to 24.58) |
| Simulated saliva | HAdV4 | 22.97 (23.40 to 22.53) |
| | H1N1pdm09 | 20.79 (18.14 to 23.44) |
| | HRV16 | 23.70 (22.10 to 25.30) |

### Supporting information figures

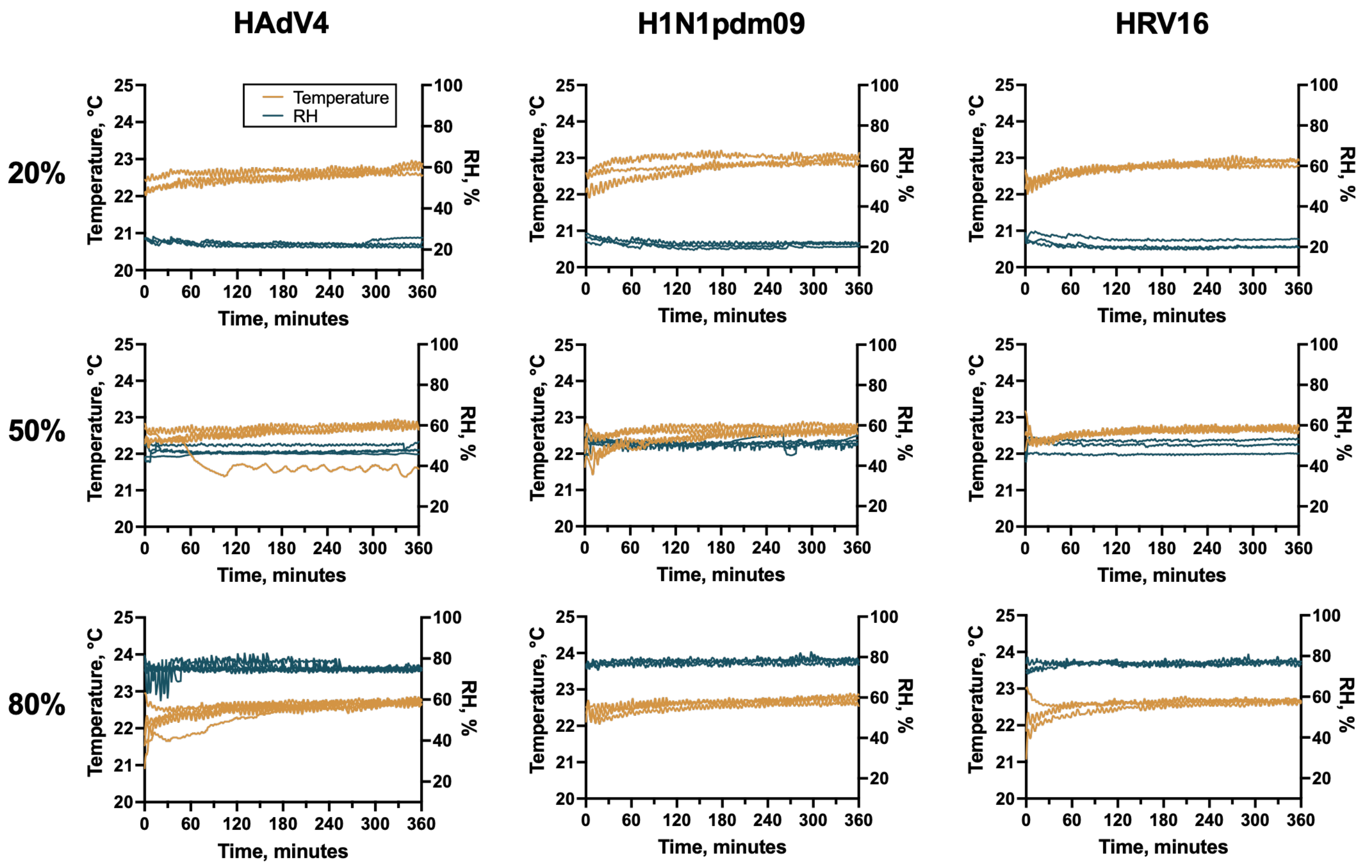

**Figure S1**. Temperature and relative humidity during human saliva deposited particle experiments. Measurements were taken every 1 min. Individual lines represent independent replicate experiments.

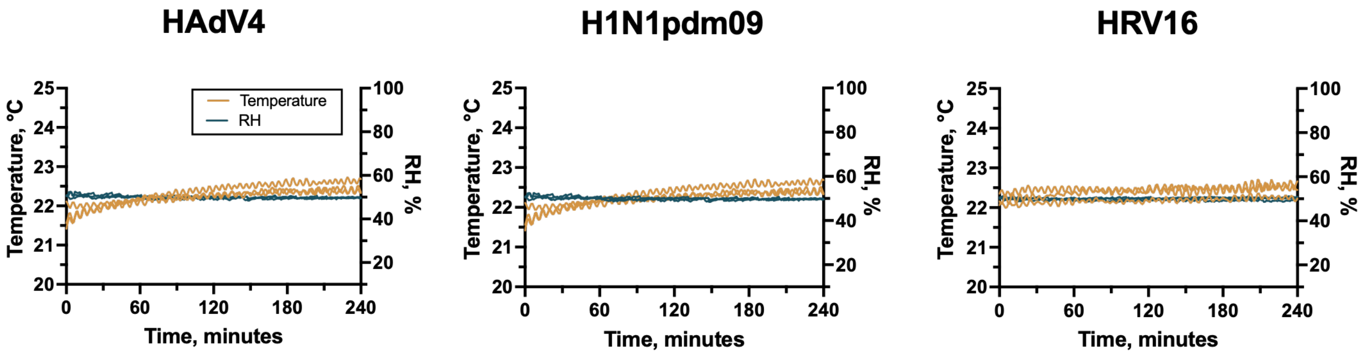

**Figure S2.** Temperature and relative humidity during simulated saliva deposited particle experiments. Measurements were taken every 1 min. Individual lines represent independent replicate experiments.

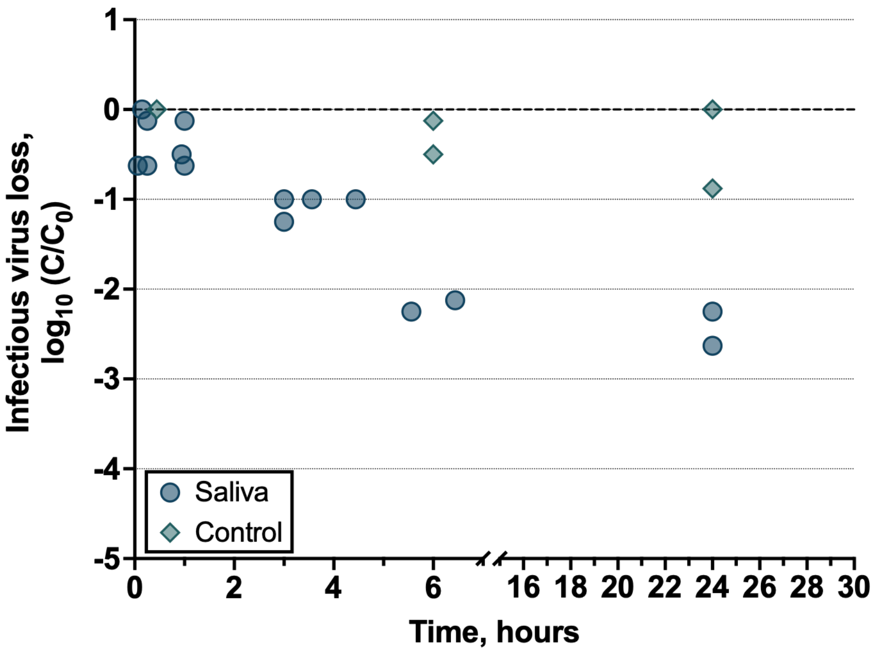

**Figure S3**. Infectious HAdV4 decay in deposited human saliva particles exposed to 50% RH at 22.5°C for 24 hours. Two technical replicates were conducted for each independent replicate, and the mean decay of two technical replicates is shown for each independent replicate (n ≥ 2). Controls are bulk solution samples taken at the beginning and end of each experiment.

.

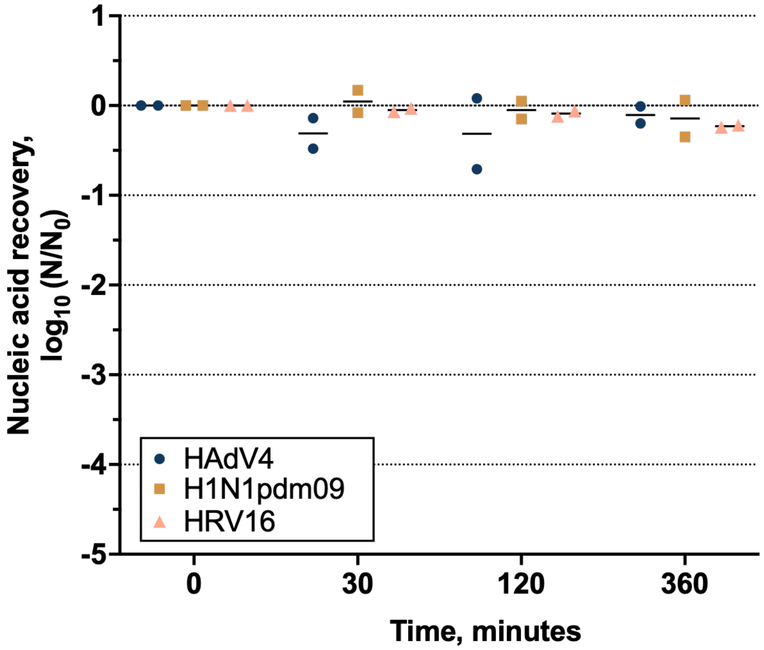

**Figure S4**. Viral nucleic acid levels recovered in virus-containing deposited saliva particles after 30 min, 2 hours, and 6 hours exposure to 50% RH at 22.5°C, for independent experiments (n = 2). Individual symbols are shown for each independent replicate, and bars indicate the mean.

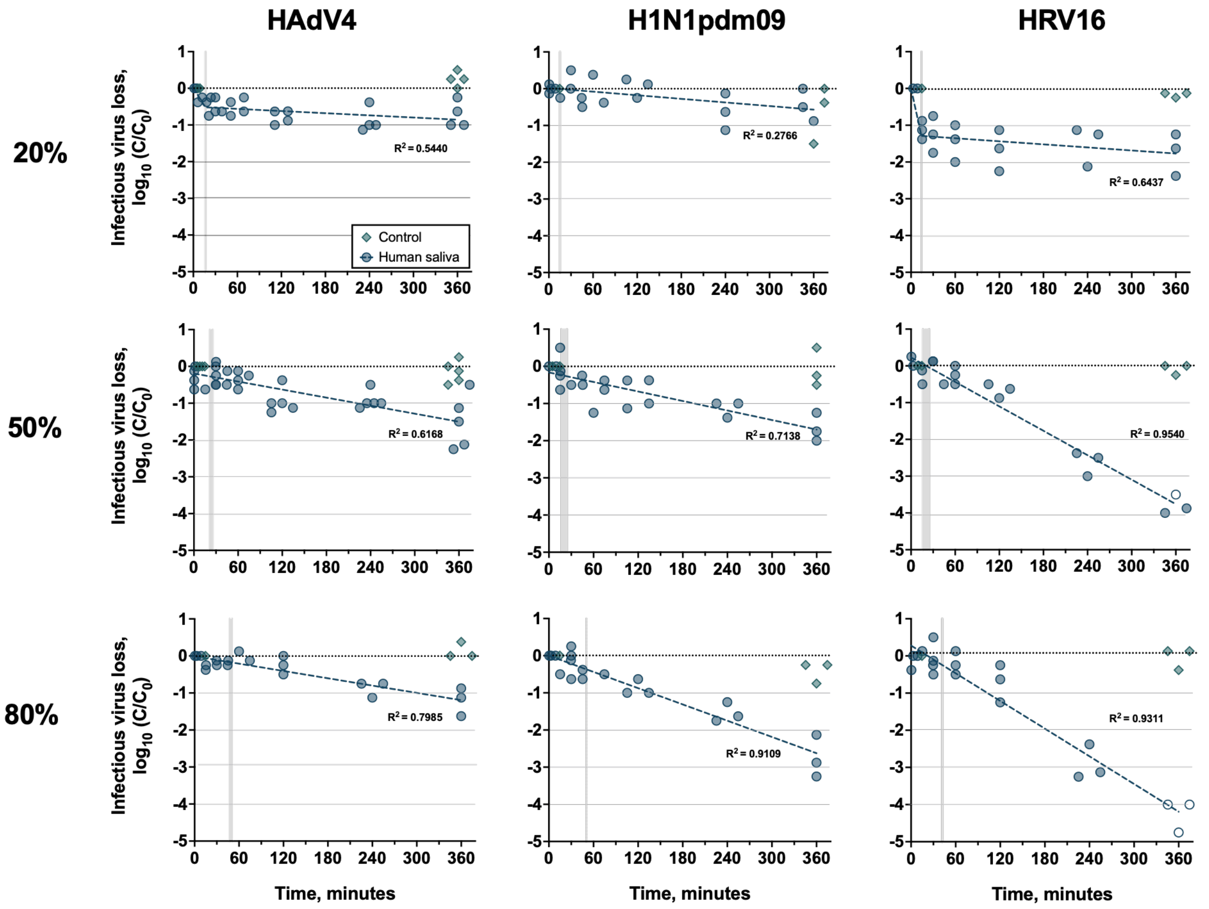

**Figure S5.** HAdV4, H1N1pdm09, and HRV16 decay kinetics in deposited human saliva particles with exposure to 20%, 50% and 80% RH fitted using the best fit linear model (single-phase or two-phase decay).

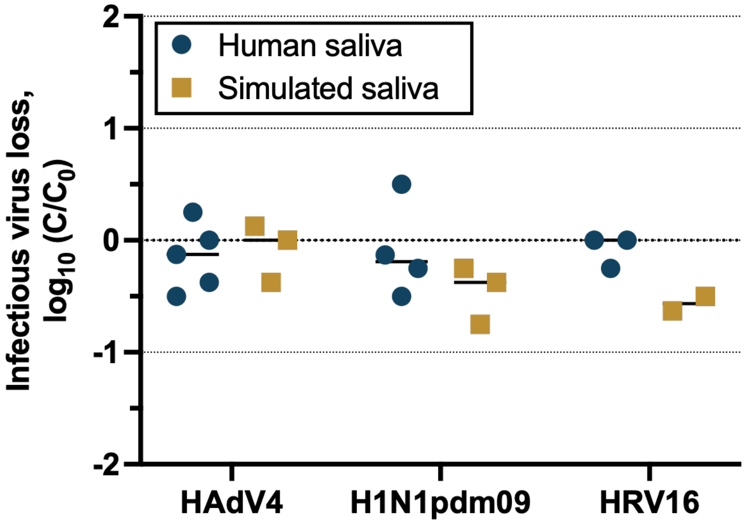

**Figure S6.** Infectious virus loss in bulk controls containing HAdV4, H1N1pdm09, or HRV16 suspended in human or simulated saliva. Symbols represent independent replicates (n ≥ 2), and lines represent mean drying times.
